# Co-translational protein targeting facilitates centrosomal recruitment of PCNT during centrosome maturation

**DOI:** 10.1101/241083

**Authors:** Guadalupe Sepulveda, Mark Antkowiak, Ingrid Brust-Mascher, Karan Mahe, Tingyoung Ou, Noemi Castro, Lana N. Christensen, Lee Cheung, Daniel Yoon, Bo Huang, Li-En Jao

**Affiliations:** Department of Cell Biology and Human Anatomy, University of California, Davis, School of Medicine, Davis, CA 95616, USA; Department of Anatomy, Physiology and Cell Biology, University of California, Davis, School of Veterinary Medicine, Davis, CA 95616, USA; Department of Pharmaceutical Chemistry, University of California, San Francisco, San Francisco, CA 94143, USA

**Author notes:** These authors contributed equally to this work (Guadalupe Sepulveda, Mark Antkowiak, Ingrid Brust-Mascher).

## Abstract

As microtubule-organizing centers of animal cells, centrosomes guide the formation of the bipolar spindle that segregates chromosomes during mitosis. At mitosis onset, centrosomes maximize microtubule-organizing activity by rapidly expanding the pericentriolar material (PCM). This process is in part driven by the large PCM protein pericentrin (PCNT), as its level increases at the PCM and helps recruit additional PCM components. However, the mechanism underlying the timely centrosomal enrichment of PCNT remains unclear. Here we show that PCNT is delivered co-translationally to centrosomes during early mitosis by cytoplasmic dynein, as evidenced by centrosomal enrichment of *PCNT* mRNA, its translation near the centrosome, and requirement of intact polysomes for *PCNT* mRNA localization. Additionally, the microtubule minus-end regulator, ASPM, is also targeted co-translationally to mitotic spindle poles. Together, these findings suggest that co-translational targeting of cytoplasmic proteins to specific subcellular destinations may be a generalized protein targeting mechanism.

## Introduction

A centrosome consists of a pair of centrioles embedded in a protein-dense matrix known as the pericentriolar material (PCM). The PCM functions as a major microtubule organizing center in animal cells (Gould & Borisy, 1977) as it serves as a platform onto which γ-tubulin ring complexes (γ-TuRCs), the main scaffold mediating microtubule nucleation, are loaded (Moritz, Braunfeld, Sedat, Alberts, & Agard, 1995; Zheng, Wong, Alberts, & Mitchison, 1995).

At the onset of mitosis, centrosomes rapidly expand their PCM. This process, termed centrosome maturation, is essential for proper spindle formation and chromosome segregation (Woodruff, Wueseke, & Hyman, 2014). Centrosome maturation is initiated by phosphorylation of core PCM components, such as Pericentrin (PCNT) and Cnn, by mitotic kinases PLK1/Polo and Aurora kinase A (Conduit, Feng, et al., 2014; Joukov, Walter, & De Nicolo, 2014; Kinoshita et al., 2005; Lee & Rhee, 2011). These events then trigger the cooperative assembly of additional PCM scaffold proteins (e.g., PCNT, CEP192/SPD-2, CEP152/Asterless, CEP215/CDK5RAP2/Cnn or SPD-5) into an expanded PCM matrix that encases the centrioles (Conduit, Richens, et al., 2014; Hamill, Severson, Carter, & Bowerman, 2002; Kemp, Kopish, Zipperlen, Ahringer, & O'Connell, 2004), culminating in the recruitment of additional γ-TuRCs and tubulin molecules that promote microtubule nucleation and render centrosomes competent for mediating the formation of bipolar spindles and chromosome segregation (Conduit, Wainman, & Raff, 2015; Gopalakrishnan et al., 2011; Woodruff et al., 2014).

Pericentrin (PCNT) is one of the first core PCM components identified to be required for spindle formation (Doxsey, Stein, Evans, Calarco, & Kirschner, 1994). Importantly, mutations in *PCNT* have been linked to several human disorders including primordial dwarfism (Anitha et al., 2009; Delaval & Doxsey, 2010; Griffith et al., 2008; Numata et al., 2009; Rauch et al., 2008). Pericentrin is an unusually large coiled-coil protein (3,336 amino acids in human) that forms elongated fibrils with its C-terminus anchored near the centriole wall and the N-terminus extended outwardly and radially across PCM zones in interphase cells (Lawo, Hasegan, Gupta, & Pelletier, 2012; Mennella et al., 2012; Sonnen, Schermelleh, Leonhardt, & Nigg, 2012). Recent studies showed that pericentrin plays an evolutionarily conserved role in mitotic PCM expansion and interphase centrosome organization, as loss of pericentrin activity in human, mice, and flies all results in failed recruitment of other PCM components to the centrosome and affects the same set of downstream orthologous proteins in each system (e.g., CEP215 in human, Cep215 in mice, and Cnn in flies) (C. T. Chen et al., 2014; Lee & Rhee, 2011; Lerit et al., 2015).

In vertebrates, a key function of PCNT is to initiate centrosome maturation (Lee & Rhee, 2011) and serve as a scaffold for the recruitment of other PCM proteins (Haren, Stearns, & Luders, 2009; Lawo et al., 2012; Purohit, Tynan, Vallee, & Doxsey, 1999; Zimmerman, Sillibourne, Rosa, & Doxsey, 2004). However, the mechanism underlying the timely synthesis and recruitment of a large sum of PCNT proteins to the PCM is as yet unresolved. Given its large size (>3,300 amino acids) and the modest rate of translation elongation (~3–10 amino acids per second, Bostrom et al., 1986; Ingolia, Lareau, & Weissman, 2011; Morisaki et al., 2016; Pichon et al., 2016; Wang, Han, Zhou, & Zhuang, 2016; Wu, Eliscovich, Yoon, & Singer, 2016; Yan, Hoek, Vale, & Tanenbaum, 2016), synthesizing a full-length PCNT protein would take ~10–20 minutes to complete after translation initiation. Notably, after the onset of mitosis, the PCM reaches its maximal size immediately before metaphase in ~30 minutes in human cells (Gavet & Pines, 2010; Lenart et al., 2007). Thus, the cell faces a kinetics challenge of synthesizing, transporting, and incorporating multiple large PCM proteins such as PCNT into mitotic centrosomes within this short time frame.

We show here that *pericentrin* mRNA is spatially enriched at the centrosome during mitosis in zebrafish embryos and cultured human cells. In cultured cells, the centrosomal enrichment of *PCNT* mRNA predominantly occurs during early mitosis, concomitantly with the peak of centrosome maturation. We further show that centrosomally localized *PCNT* mRNA undergoes active translation and that acute inhibition of translation compromises the incorporation of PCNT proteins into the centrosome during early mitosis. Moreover, we find that centrosomal localization of *PCNT* mRNA requires intact polysomes, microtubules, and cytoplasmic dynein activity. Taken together, our results support a model in which translating *PCNT* polysomes are being actively transported toward the centrosome during centrosome maturation. We propose that by targeting actively translating polysomes toward centrosomes, the cell can overcome the kinetics challenge of synthesizing, transporting, and incorporating the unusually large PCNT proteins into the centrosome. Lastly, we find that the cell appears to use a similar co-translational targeting mechanism to synthesize and deliver another unusually large protein, the microtubule minus-end regulator, ASPM, to the mitotic spindle poles. Thus, co-translational protein targeting might be a mechanism widely employed by the cell to transport cytoplasmic proteins to specific subcellular compartments and organelles.

## Results

### Zebrafish *pcnt* mRNA is localized to the centrosome in blastula-stage embryos

We found that *pericentrin* (*pcnt*) transcripts were localized to distinct foci in early zebrafish embryos, whereas those of three other core PCM components, *cep152, cep192*, and *cep215*, showed a pan-cellular distribution (Figure 1A). This striking *pcnt* mRNA localization was observed using two independent, non-overlapping antisense probes against the 5’ or 3’ portion of RNA (Figure 1B). The specificity of *in situ* hybridization was further confirmed by the loss of signals in two frameshift maternal-zygotic *pcnt* knockout embryos (MZ*pcnt^tup2^* and MZ*pcnt^tup5^*) (Figure 1B and Figure 1-figure supplement 1), where the *pcnt* transcripts were susceptible to nonsense-mediated decay pathway. By co-staining with the centrosome marker γ-tubulin, we demonstrated that zebrafish *pcnt* mRNA is specifically localized to the centrosome (Figure 1C).

**Figure 1.**
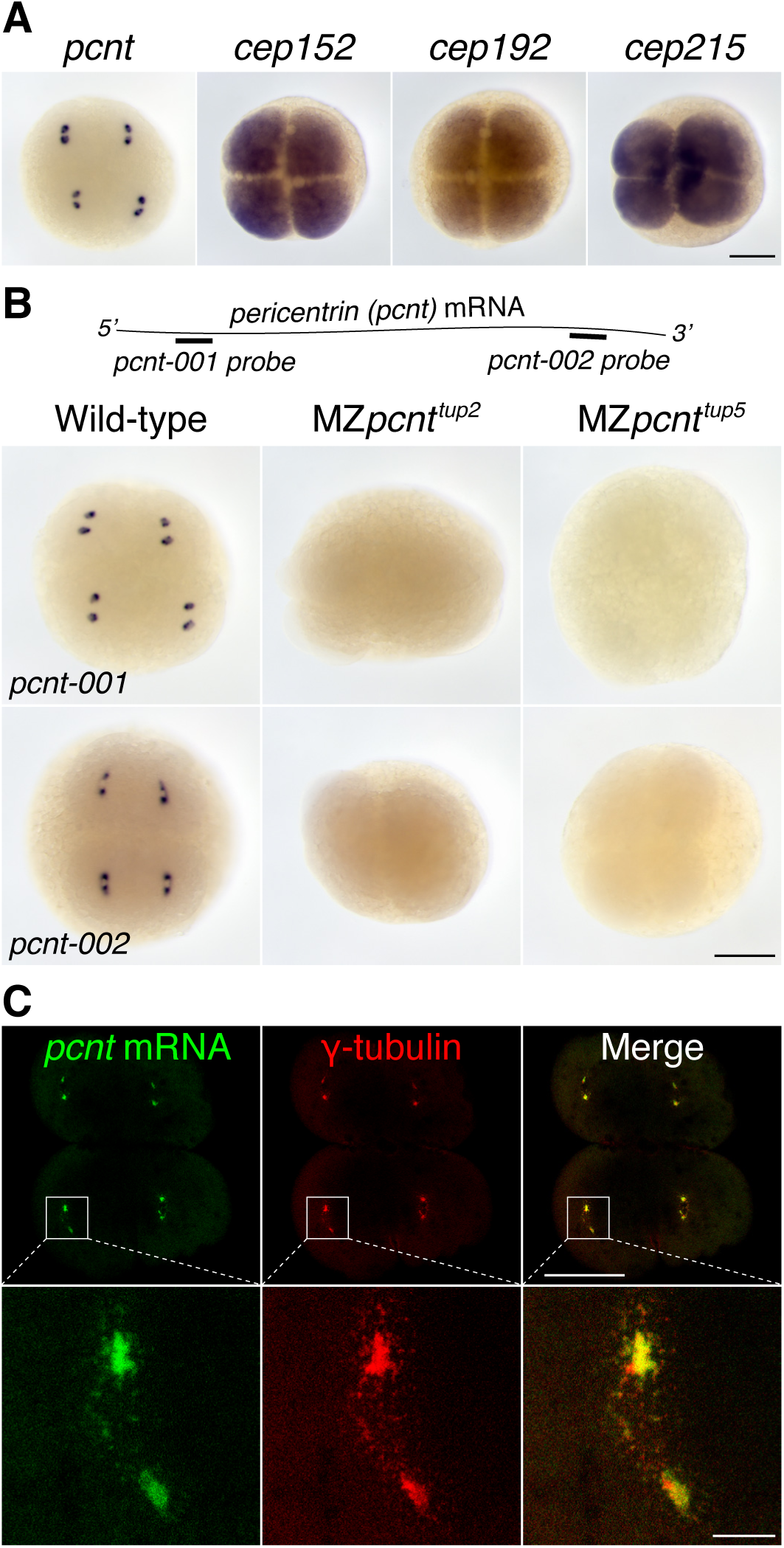
*Pericentrin (pcnt)* mRNA is localized to centro-somes in early zebrafish embryos. **(A)** RNA *in situ* hybridization of transcripts of different PCM components in 4-cell stage zebrafish embryos. Note that while the mRNA of *cep152, cep192*, and *cep215* displayed a pan-cellular distribution, *pcnt* mRNA was concentrated at two distinct foci in each cell. **(B)** RNA *in situ* hybridization showed similar dot-like patterns of *pcnt* transcripts with two non-overlapping antisense probes. The signals were lost in two maternal-zygotic (MZ) *pcnt* mutants. **(C)** Fluorescent RNA *in situ* hybridization and anti-γ-tubulin co-staining demonstrated the centrosomal localization of *pcnt* mRNA. Scale bars: 200 *μ*m or 25 *μ*m (inset in **C**).

### Human *PCNT* mRNA is enriched at the centrosome during early mitosis

To test whether centrosomal localization of *pcnt* mRNA is conserved beyond early zebrafish embryos, we examined the localization of human *PCNT* mRNA in cultured HeLa cells using fluorescent *in situ* hybridization (FISH). Consistent with our observation in zebrafish, human *PCNT* mRNA was also localized to the centrosome (Figure 2). Interestingly, this centrosomal enrichment of *PCNT* mRNA was most prominent during early mitosis (i.e., prophase and prometaphase) and declined after prometaphase. The signal specificity was confirmed by two non-overlapping probes against the 5’ or 3’ portion of the *PCNT* transcript (Figure 2-figure supplement 1A). Furthermore, using an alternative FISH method, Stellaris^®^ single-molecule FISH (smFISH) against the 5’ or 3’ portion of the *PCNT* transcript, we observed highly similar centrosomal enrichment of *PCNT* mRNA during early mitosis, with near single-molecule resolution (Figure 2-figure supplement 1B). Similar smFISH results were observed in both HeLa and RPE-1 cells (data not shown). Together, these results indicate that *PCNT* mRNA is specifically enriched at the centrosome during early mitosis in cultured human cells. We speculate that the seemingly constant presence of zebrafish *pcnt* mRNA at the centrosome of early blastula-stage embryos is due to the fast cell cycle without gap phases at this stage (~20 minutes per cycle).

**Figure 2.**
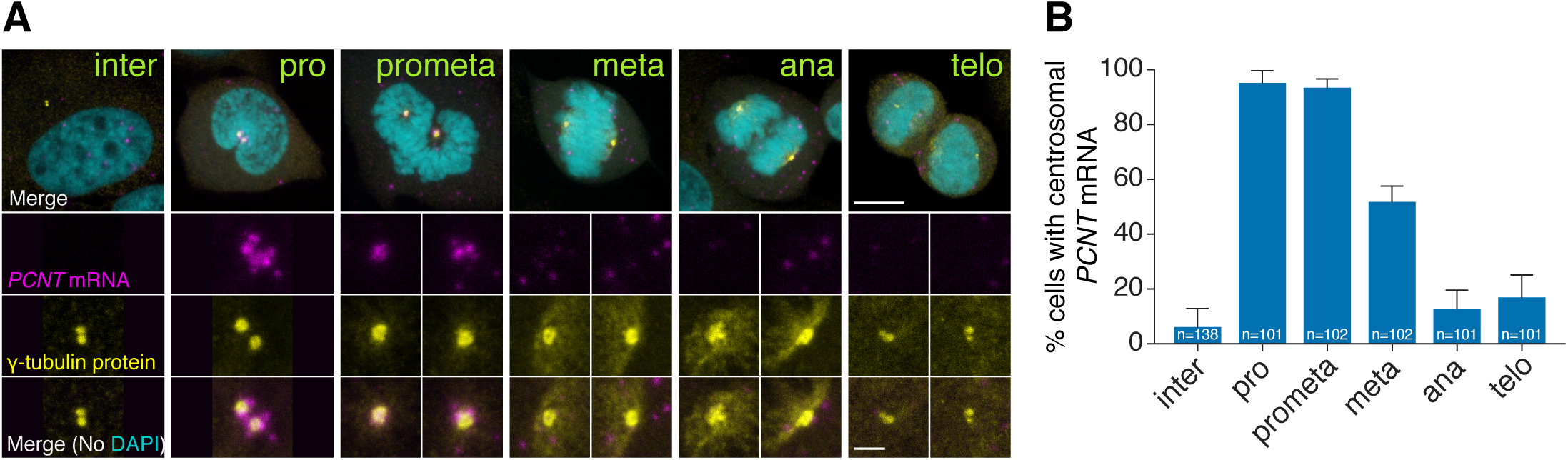
Human *PCNT* mRNA is localized to centrosomes during early mitosis. **(A)** Synchronized HeLa cells were subjected to fluorescent *in situ* hybridization with tyramide signal amplification against *PCNT* mRNA and anti-γ-tubulin immunostaining. Note that *PCNT* mRNA was localized to centrosomes predominantly during prophase (pro) and prometaphase (prometa). **(B)** Quantification of *PCNT* mRNA localization at centrosomes during cell cycle stages from three experimental replicates. Data are represented as mean with standard deviation (SD) with the total number of cells analyzed indicated. Scale bars: 10 *μ*m and 2 *μ*m (inset).

### Zebrafish *pcnt* mRNA is localized to the centrosome of mitotic retinal neuroepithelial cells *in vivo*

We next tested whether centrosomal localization of *pcnt* mRNA also takes place in differentiated tissues *in vivo*. We focused on the retinal neuroepithelia of 1 day old zebrafish because at this developmental stage, retinal neuroepithelial cells in different cell cycle stages can be readily identified based on the known patterns of interkinetic nuclear migration (e.g., mitotic cells at the apical side of retina) (Baye & Link, 2007). Again, we observed that zebrafish *pcnt* mRNA was enriched at the centrosome of mitotic, but not of non-mitotic, neuroepithelial cells (Figure 2-figure supplement 2). We thus conclude that centrosomal enrichment of *pericentrin* mRNA is likely a conserved process in mitotic cells.

### Centrosomally localized *PCNT* mRNA undergoes active translation

Interestingly, the timing of this unique centrosomal accumulation of *PCNT* mRNA in cultured cells (Figure 2) overlaps precisely with that of centrosome maturation (Khodjakov & Rieder, 1999; Piehl, Tulu, Wadsworth, & Cassimeris, 2004). These observations raise the intriguing possibility that *PCNT* mRNA might be translated near the centrosome to facilitate the incorporation of PCNT proteins into the PCM during centrosome maturation.

To determine whether *PCNT* mRNA is actively translated near the centrosome, we developed a strategy to detect actively translating PCNT polysomes by combining *PCNT* smFISH and double immunofluorescence to label *PCNT* mRNA, and the N- and C-termini of PCNT protein simultaneously (Figure 3A). Given the inter-ribosome distance of approximately 260 nucleotides on a transcript during translation (Wang et al., 2016) and the large size of *PCNT* mRNA (10 knt), a single *PCNT* transcript can be actively translated by as many as 40 ribosomes simultaneously. Therefore, up to 40 nascent polypeptides emerging from a single *PCNT* polysome can be visualized by anti-PCNT N-terminus immunostaining. By combining this immunostaining strategy with *PCNT* smFISH, multiple nascent PCNT polypeptides can be visualized on a single *PCNT* mRNA. Furthermore, the signals from antibody staining are determined by the location of the epitopes. Therefore, the translating nascent PCNT polypeptides, with the C-terminus not yet synthesized, would only show positive signals from anti-PCNT N-terminus immunostaining (and be positive for *PCNT* smFISH), whereas fully synthesized PCNT protein would show signals from both anti-PCNT N- and C-terminus immunostaining (and be negative for *PCNT* smFISH because of release of the full-length protein from the RNA-bound polysomes).

**Figure 3.**
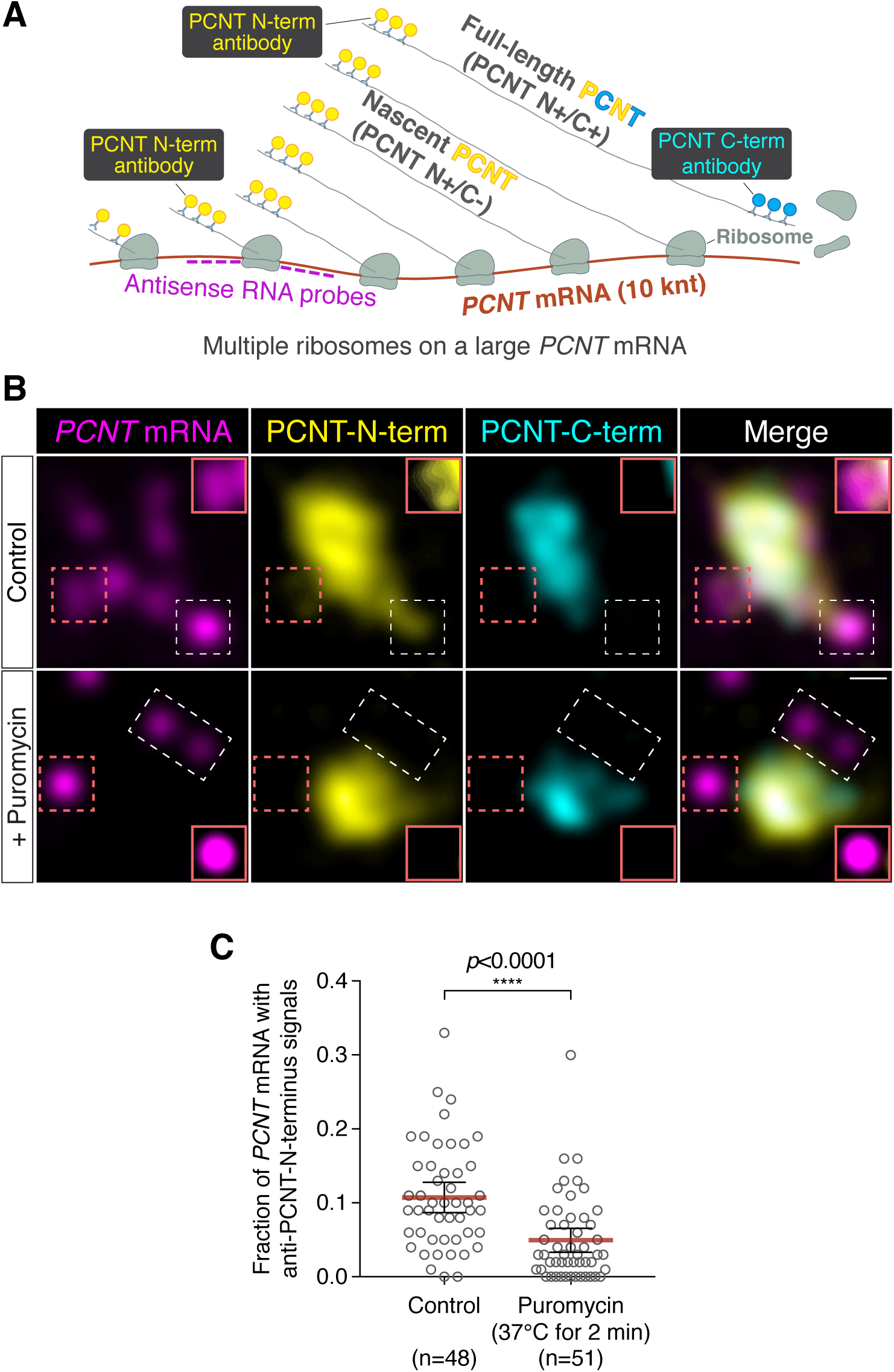
Centrosomally localized *PCNT* mRNA undergoes active translation. **(A)** A strategy of using smFISH and double immunofluorescence (IF) to distinguish between newly synthesized and full-length PCNT proteins. **(B)** Prometaphase HeLa cells were subjected to *PCNT* smFISH and anti-PCNT immunostaining against the N- and C-terminus of PCNT protein (PCNT-N-term and PCNT-C-term). Note that the putative active translation sites were labeled by PCNT-N-term IF and *PCNT* smFISH, but not by PCNT-C-term IF (top row). However, upon the puromycin treatment (300 *μ*m for 2 minutes at 37°C, bottom row), PCNT-N-term IF signals were no longer colocalized with *PCNT* smFISH signals, indicating that those PCNT-N-term IF signals on RNA represent nascent PCNT polypeptides. Orange boxes show higher contrast of selected areas (dashed orange boxes) for better visualization. **(C)** *PCNT* smFISH signals between 1 and 3 *μ*m radius from the centrosome center were quantified for the presence of anti-PCNT-N-term IF signals with or without a short puromycin treatment. Data are represented as mean ± 95% CI (confidence intervals), with the number of cells analyzed indicated. *p* value was obtained with Student's t-test (two-tailed). Scale bar: 0.5 *μ*m.

Using this strategy, we detected nascent PCNT polypeptides emerging from *PCNT* mRNA near the centrosome during early mitosis (Figure 3B, top row, PCNT N^+^/C^-^/*PCNT* smFISH^+^). As an important control, we showed that colocalization of *PCNT* mRNA with anti-PCNT N-terminus signals was lost after a brief treatment of cells with puromycin (Figure 3B, bottom row), under a condition confirmed to inhibit translation by dissociating the ribosomes and releasing the nascent polypeptides (Figure 3-figure supplement 1, Wang et al., 2016; Yan et al., 2016). Next, we developed a methodology to quantify the effect of puromycin treatment on the colocalization of *PCNT* mRNA and anti-PCNT N-terminus signals in three dimensional (3D) voxels rendered from confocal z-stacks. Given that the mean radius of a mitotic centrosome is ~1 μm (Figure 3-figure supplement 2), we specifically quantified the fraction of *PCNT* mRNA between 1 and 3 μm from the center of each centrosome—i.e., the RNA close to, but not within, the centrosome—with anti-PCNT N-terminus signals in early mitotic cells, with or without the brief puromycin treatment. Consistent with the results shown in Figure 3B, upon the short puromycin treatment, the fraction of *PCNT* mRNA with anti-PCNT N-terminus signals was significantly reduced, with many *PCNT* mRNA no longer bearing anti-PCNT N-terminus signals (Figure 3C). Together, these results indicate that during early mitosis, a population of *PCNT* mRNA is undergoing active translation near the centrosome.

### Centrosomal localization of *pcnt/PCNT* mRNA requires intact polysomes, microtubules, and dynein activity

In addition to the loss of anti-PCNT N-terminus signals from *PCNT* mRNA, surprisingly, the brief puromycin treatment led to the population of *PCNT* mRNA shifting away from the centrosome (Figure 4A and B). Similarly, when zebrafish embryos were injected with puromycin at the 1-cell stage, *pcnt* transcripts became diffused throughout the cell (Figure 4-figure supplement 1). Because puromycin dissociates ribosomes and nascent polypeptides, these observations suggest that *PCNT/pcnt* mRNAs in human and zebrafish are enriched near the centrosome by tethering to the actively translating ribosomes.

**Figure 4.**
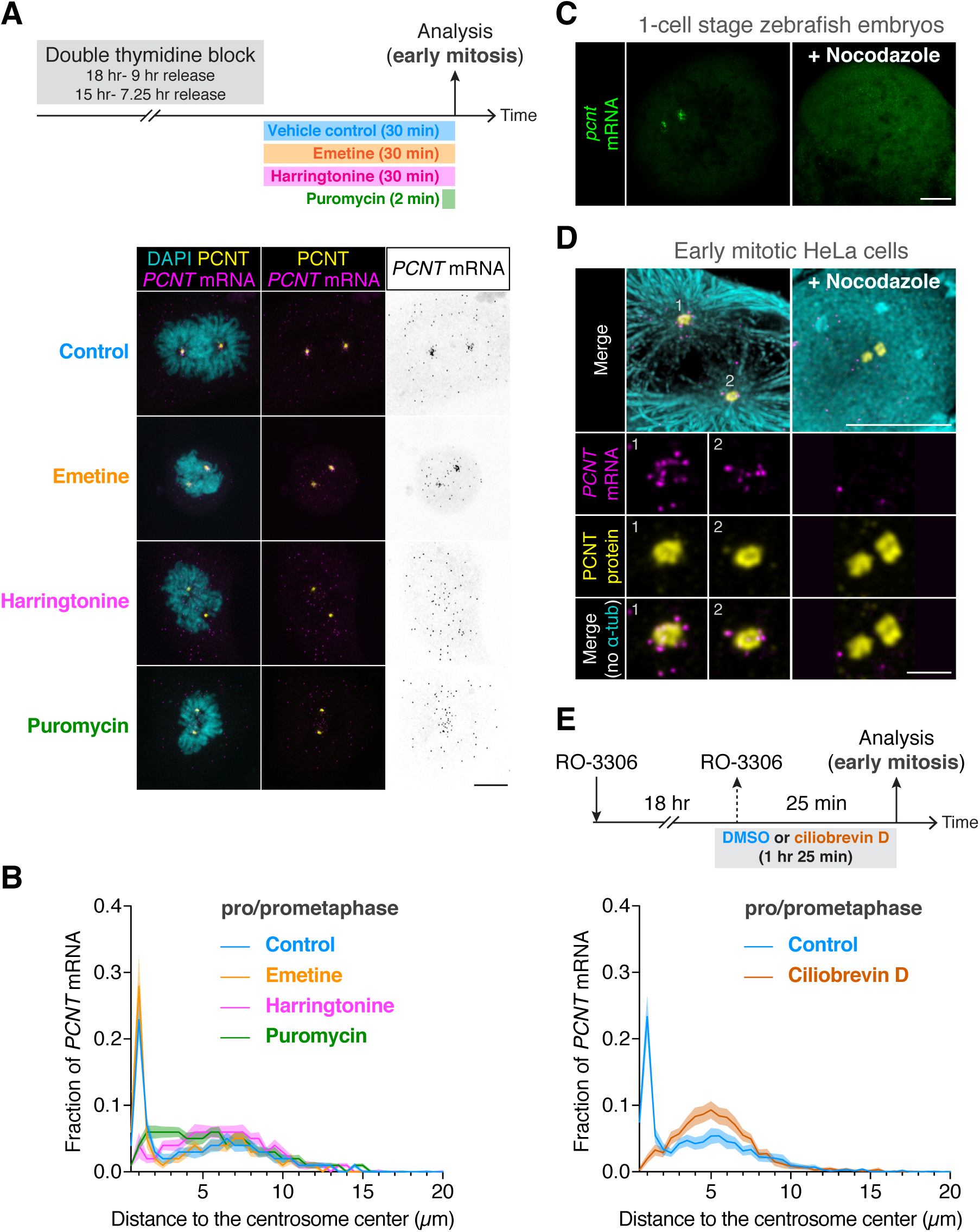
Centrosomal localization of *pcnt/PCNT* mRNA requires intact polysomes, microtubules, and dynein activity. **(A)** HeLa cells were synchronized by a double thymidine block and treated with DMSO vehicle (Control), 208 *μ*m emetine, 3.76 *μ*m harringtonine for 30 minutes, or 300 *μ*m puromycin for 2 minutes before anti-PCNT immunostaining and *PCNT* smFISH. Representative confocal images are shown for each condition. **(B)** The distribution of *PCNT* mRNA in cells was quantified by measuring the distance between 3D rendered *PCNT* smFISH signals and the center of the nearest centrosome (labeled by anti-PCNT immunostaining). The fractions of mRNA as a function of distance to the nearest centrosome (binned in 0.5 *μ*m intervals) were then plotted as mean (solid lines) ± 95% CI (shading). Note that *PCNT* mRNA moved away from the centrosome upon the puromycin or harringtonine treatment, but stayed close to the centrosome upon the emetine treatment, similar to the control. n=45, 48, 57, and 51 for control, emetine, puromycin, and harringtonine conditions, respectively, from three independent experiments. **(C)** Zebrafish embryos were injected with DMSO vehicle or 100 *μ*g/ml nocodazole at the 1-cell stage followed by *pcnt* FISH. **(D)** HeLa cells were treated with DMSO vehicle or 3 *μ*g/ml nocodazole for 2 hours at 37°C before anti-α-tubulin, anti-PCNT immunostaining, and *PCNT* smFISH. Note that *pcnt/PCNT* mRNA in early embryos **(C)** and in early mitotic cells **(D)** was no longer enriched at the centrosome after microtubules were depolymerized. **(E)** HeLa cells were synchronized by RO-3306 and treated with DMSO vehicle or 50 *μ*m ciliobrevin D for 1 hour 25 minutes before anti-PCNT immunostaining and *PCNT* smFISH. The distribution of *PCNT* mRNA in cells was quantified as in **(B)**. n=70 and 63 for control and ciliobrevin D conditions, respectively, from a representative experiment (two technical duplicates per condition). Note that *PCNT* mRNA was no longer enriched at the centrosome upon the ciliobrevin D treatment. Scale bars, 10 *μ*m **(A)**, 100 *μ*m **(C)**, 10 *μ*m **(D)**, and 2 *μ*m (inset in **D**).

To further test the dependency of centrosomal enrichment of *PCNT* mRNA on intact, actively translating polysomes, we treated the cultured cells with either emetine, which stabilizes polysomes by irreversibly binding the ribosomal 40S subunit and thus “freezing” translation during elongation (Jimenez, Carrasco, & Vazquez, 1977), or harringtonine, which disrupts polysomes by blocking the initiation step of translation while allowing downstream ribosomes to run off from the mRNA (Huang, 1975). We found that *PCNT* mRNA localization patterns in emetine- and harringtonine-treated cells resembled those observed in vehicle- (control) and puromycin-treated cells, respectively (Figure 4A and B). Congruent with the detection of nascent PCNT polypeptides near the centrosome (Figure 3), these data support the model that centrosomal enrichment of *PCNT* mRNA relies on centrosomal enrichment of polysomes that are translating *PCNT* mRNA.

We often observed that the two centrosomes in early mitotic cells were asymmetric in size where more *PCNT* mRNA was enriched near the larger centrosome (Figure 4-figure supplement 2). Because the microtubule nucleation activity is often positively correlated with the centrosome size, we speculated that centrosomal enrichment of *pericentrin* mRNA/polysomes might be a microtubule-dependent process. We thus tested if the localization of *pericentrin* mRNA would be perturbed when microtubules were depolymerized. We found that in both zebrafish and cultured human cells, *pcnt/PCNT* mRNA was no longer enriched around the centrosome upon microtubule depolymerization (Figure 4C and D). In contrast, a cytochalasin B treatment, which disrupts the actin cytoskeleton, had no effect on the centrosomal enrichment of *PCNT* mRNA (Figure 4-figure supplement 3). These results suggest that microtubules, but not actin filaments, serve as “tracks” on which *pericentrin* mRNA/polysomes are transported.

Given that cytoplasmic dynein is a common minus-end-directed, microtubule-based motor that transports cargo toward the microtubule minus end (i.e., toward the centrosome), we next tested whether centrosomal localization of *PCNT* mRNA is a dynein-dependent process. We treated the cells with ciliobrevin D, a specific small molecule inhibitor of cytoplasmic dynein (Firestone et al., 2012) and quantified the effect of this treatment on the centrosomal localization of *PCNT* mRNA. We found that *PCNT* mRNA was no longer enriched at the centrosome upon the ciliobrevin D treatment (Figure 4E). Together, these results indicate that centrosomal enrichment of *pericentrin* mRNA during early mitosis is a translation-, microtubule- and dynein-dependent process.

### Active translation of *PCNT* mRNA during early mitosis contributes to the optimal incorporation of PCNT protein into the mitotic PCM

To determine the functional significance of translation of centrosomally localized *PCNT* mRNA during early mitosis, we compared centrosomal PCNT levels shortly before and after mitotic entry (i.e., late G2 vs. early M phase). We arrested cultured human cells from progression out of late G2 phase using the CDK1 inhibitor RO-3306 (Vassilev et al., 2006). CDK1 is largely inactive during G2 and becomes activated at the onset of mitosis (Gavet & Pines, 2010; M. Jackman, Lindon, Nigg, & Pines, 2003). In the presence of RO-3306, cells can be held at late G2 phase, and upon inhibitor washout, cells can be released into mitosis. Because cell cycle synchronization is rarely 100% homogeneous in a cell population, we decided to quantify the amount of centrosomal PCNT at the single cell level using anti-PCNT immunostaining of individual cells. To confidently identify late G2 cells in RO-3306-treated population, we used a RPE-1 cell line stably expressing Centrin-GFP (Uetake et al., 2007) and categorized the cells as “late G2” if (1) their two centrosomes (with two centrin dots per centrosome) were separated by > 2 *μ*m—a sign indicating the loss of centrosome cohesion that occurs during late G2 to M transition (Bahe, Stierhof, Wilkinson, Leiss, & Nigg, 2005; Fry et al., 1998; Mardin, Agircan, Lange, & Schiebel, 2011) and (2) their DNA was not condensed. We identified the cells as early M phase cells (i.e., prophase or prometaphase) 25 minutes after RO-3306 washout by observing DNA morphology.

Using this strategy, we found that approximately 2-fold more PCNT proteins were incorporated into the centrosomes in early mitotic cells as compared to late G2 cells (Figure 5A). Importantly, the numbers of *PCNT* mRNA did not significantly differ between late G2 and early M phases, even though there was an approximately 4-fold increase from G1 to late G2 phases (Figure 5B). Therefore, these results indicate that the increase in centrosomal PCNT protein levels when cells progress from G2 to M phases (e.g., the 25-minute period after RO-3306 washout) is due to upregulation of translation and not to altered mRNA abundance.

**Figure 5.**
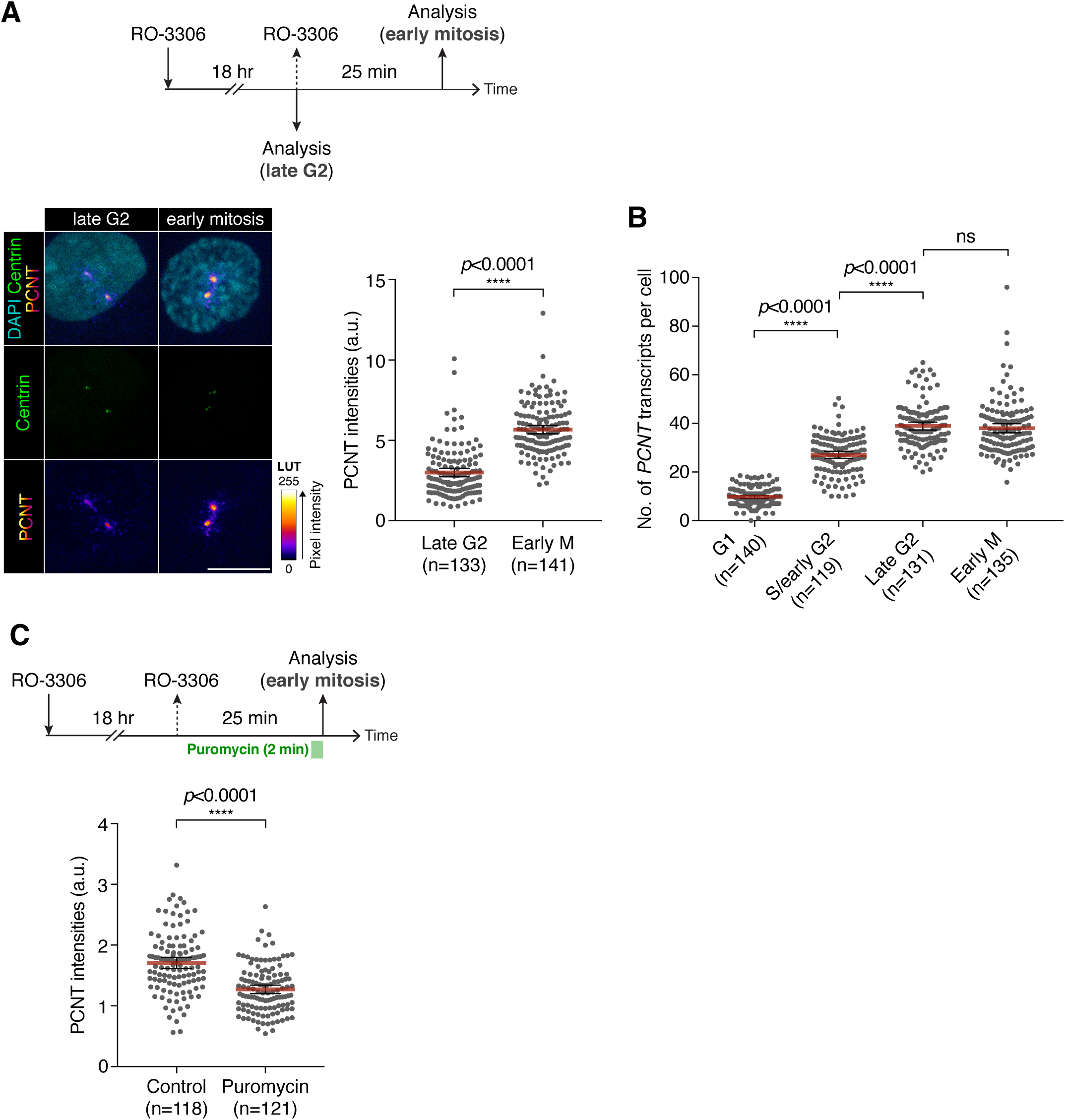
Centrosomal localization of *PCNT* mRNA/polysomes contributes to PCNT incorporation into mitotic centrosomes. **(A)** Centrin-GFP RPE-1 cells—at either late G2 or early M phase—were subjected to anti-PCNT immunostaining. Representative confocal images are shown for each condition. A “fire” lookup table (LUT) was used to show PCNT signal intensities. The sum intensity of anti-PCNT signals from both centrosomes of each cell was measured and plotted. **(B)** Numbers of *PCNT* mRNA at different cell cycle stages of Centrin-GFP RPE-1 cells were determined by *PCNT* smFISH. S phase/early G2 cells were identified by EdU labeling for 30 minutes. **(C)** HeLa cells were treated with vehicle control or 300 *μ*Μ puromycin for 2 minutes before anti-PCNT immunostaining. The sum intensity of anti-PCNT signals from both centrosomes of each prophase or prometaphase cell was measured and plotted. Data are represented as mean ± 95% CI. “n” indicates the total number of cells analyzed from at least two independent experiments. *p* values were obtained with Student's t-test (two-tailed). Scale bar: 10 *μ*m.

To independently assess the impact of translation during early mitosis on PCNT incorporation into the centrosomes, we disrupted this process by pulsing the RO-3306 synchronized cells with puromycin to inhibit translation for two minutes, followed by immediate fixation and anti-PCNT immunostaining. As previously shown, this condition inhibits translation acutely and dissociates PCNT nascent polypeptides from *PCNT* mRNA-containing polysomes, including those near the centrosome (Figure 3). We found that in the puromycin-treated cells, ~30% fewer PCNT molecules were incorporated into the PCM than in the control cells during prophase/prometaphase (Figure 5C). These results indicate that active translation during prophase/prometaphase is required for efficient incorporation of PCNT into the mitotic centrosomes; disruption of this process, even just briefly, significantly affects the PCNT level at the centrosomes.

Collectively, these results indicate that active translation of *PCNT* mRNA during early mitosis contributes to the optimal incorporation of PCNT proteins into the mitotic PCM and that this is most plausibly achieved by co-translational targeting of the *PCNT* mRNA-containing polysomes to the proximity of the mitotic centrosomes.

### *ASPM* mRNA is enriched at the centrosome in a translation-dependent manner during mitosis

To determine if the cell uses a similar co-translational targeting strategy to target other large proteins to the centrosome, we examined the distribution of *CEP192, CDK5RAP2/CEP215*, and *ASPM* mRNA in cultured human cells. We found that while *CEP192* and *CEP215* mRNA did not show any centrosomal enrichment during early mitosis (data not shown), *ASPM* mRNA was strongly enriched at the centrosome during prometaphase and metaphase in both HeLa and RPE-1 cells (Figure 6). Furthermore, upon a short puromycin treatment, *ASPM* mRNA became dispersed throughout the cell, indicating that centrosomal enrichment of *ASPM* mRNA also requires intact polysomes as in the case with *PCNT* mRNA. ASPM (and its fly ortholog Asp) is not a PCM component *per se*, but a microtubule minus-end regulator (Jiang et al., 2017) and a spindle-pole focusing factor (Ito & Goshima, 2015; Ripoll, Pimpinelli, Valdivia, & Avila, 1985; Tungadi, Ito, Kiyomitsu, & Goshima, 2017). It is highly enriched at the mitotic spindle poles, particularly from early prometaphase to metaphase (Ito & Goshima, 2015; Jiang et al., 2017; Tungadi et al., 2017). Therefore, these data demonstrate another example of spatiotemporal coupling between active translation and translocation of polysomes to the final destination of the protein being synthesized.

**Figure 6.**
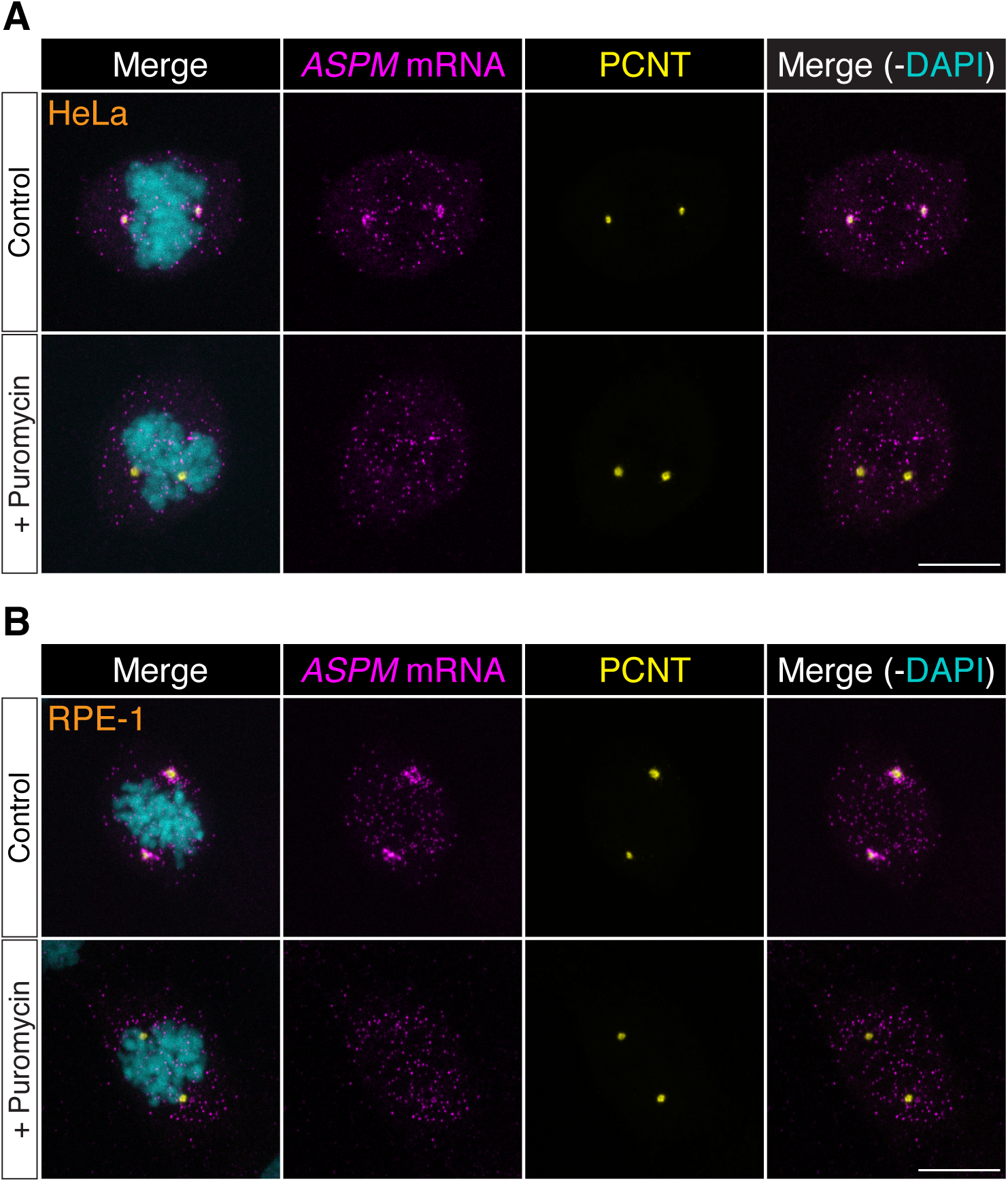
*ASPM* mRNA is enriched at centrosomes in a translation-dependent manner during mitosis. Prometaphase HeLa **(A)** or RPE-1 **(B)** cells were treated with vehicle (Control) or 300 puromycin (+ Puromycin) for 2 minutes at 37°C before fixation. The cells were subjected to *ASPM* smFISH and anti-PCNT immunostaining. Note that *ASPM* mRNA was enriched at the centrosomes/spindle poles of the prometaphase cells, but became dispersed throughout the cell upon a short puromycin treatment. Scale bars: 10 *μ*m.

## Discussion

Here we report that PCNT protein is delivered co-translationally to the centrosome during centrosome maturation through a microtubule- and dynein-dependent process. This process is demonstrated by centrosomal enrichment of *PCNT* mRNA, its translation near the centrosome, and requirement of intact translation machinery for *PCNT* mRNA localization during early mitosis. The translation- and microtubule-dependent centrosomal enrichment of *pericentrin* mRNA is observed in both zebrafish embryos and human somatic cell lines. Interestingly, the mRNA of the sole *pcnt* ortholog, *plp*, of *Drosophila melanogaster*, was also previously reported to localize to the centrosome in early fly embryos (Lecuyer et al., 2007). Although it has not been shown if the centrosomally localized *plp* mRNA undergoes active translation, it is tempting to speculate that co-translational targeting of PCNT (and its orthologous proteins) to the centrosome is an evolutionarily conserved process. In addition to PCNT, the cell appears to use a similar co-translational targeting strategy to deliver the large microtubule minus-end regulator/spindle-pole focusing factor, ASPM, to mitotic spindle poles, as *ASPM* mRNA is strongly enriched at mitotic spindle poles in a translation-dependent manner, concomitantly with the ASPM protein level reaching its maximum at the same place. We suspect that co-translational targeting of polysomes translating a subset of cytoplasmic proteins to specific subcellular destinations is a widespread mechanism used in post-transcriptional gene regulation.

### Evidence supporting translation of *PCNT* mRNA near the centrosome

In this study, we also developed a strategy of visualizing active translation. We took advantage of the large size of *PCNT* mRNA and combined *PCNT* smFISH and immunofluorescence against the N- or C-terminal epitopes of PCNT nascent polypeptides to detect which *PCNT* mRNA molecules were undergoing active translation (Figure 3). This imaging-based method allowed us to determine whether the PCNT was being newly synthesized “on site” or the PCNT was made somewhere within the cell and then transported/diffused to the centrosome because only the former would show positive signals for N-, but not C-terminus immunostaining of the synthesized protein, and these signals would be sensitive to the puromycin treatment. However, detecting nascent PCNT polypeptides by anti-PCNT N-terminus antibody staining relies on multiple copies of polypeptides tethered to the translating ribosomes for generating detectable fluorescent signals. Therefore, this method is biased toward detecting the translating *PCNT* polysomes at later stages of translation elongation, when multiple ribosomes have been loaded and multiple copies of PCNT polypeptides are available for antibody detection. This method, however, would likely fail to detect anti-PCNT N-terminus signals on the mRNA that just started to be translated. We speculate that this could explain why not all centrosomally localized *PCNT* mRNAs showed anti-PCNT N-terminus signals, although most of these *PCNT* mRNAs would shift away from the centrosome upon the puromycin or harringtonine treatment (Figure 4).

### Significance of co-translational targeting of PCNT to the centrosome during mitosis

What might be the biological significance of co-translational targeting of unusually large proteins such as PCNT or ASPM to the centrosome during mitosis? In the case of PCNT, we propose three possible mutually inclusive reasons. First, since PCNT has been placed upstream as a scaffold to initiate centrosome maturation (Lee & Rhee, 2011) and to help recruit other PCM components, including NEDD1, CEP192, and CDK5RAP2 (Lawo et al., 2012), it is critical to have optimal amounts of PCNT incorporated at the centrosome early during mitosis. As a polysome can synthesize multiple copies of polypeptides from a single mRNA template, mechanistically coupling translation and translocation of polysomes toward the destination of the protein being synthesized can maximize efficiency of protein production and delivery, especially for large proteins such as PCNT, which requires 10–20 minutes to synthesize. Therefore, using this co-translational targeting mechanism can enable the cell to overcome the kinetics challenge of generating and incorporating the unusually large PCNT to the centrosome efficiently before metaphase. Second, generating PCNT proteins elsewhere in the cell might be deleterious. For example, non-centrosomal accumulation of PCNT might recruit other PCM components to the unwanted locations, resulting in ectopic PCM assembly; co-translational targeting of PCNT on defined microtubule tracks through the dynein motor can help confine most full-length PCNT proteins to the centrosome. Third, co-translational targeting of nascent PCNT polypeptides might be an integrated part of mitotic PCM expansion. Akin to the co-translational targeting of membrane and secreted proteins to the endoplasmic reticulum (ER), where the translating nascent polypeptides undergo protein folding and post-translational modifications in the ER lumen (Bergman & Kuehl, 1979; W. Chen, Helenius, Braakman, & Helenius, 1995), co-translational targeting of nascent PCNT polypeptides might promote their proper folding and complex formation near the PCM, thereby facilitating integration into the expanding PCM during early mitosis. Future experiments specifically perturbing this co-translational targeting process should help distinguish these hypotheses.

### Mechanism of co-translational targeting

How are the polysomes actively translating PCNT or ASPM transported to the centrosome? In the case of PCNT, previous studies have shown that PCNT protein is transported to the centrosome through its interaction with cytoplasmic dynein (Purohit et al., 1999; Young et al., 2000), specifically through the dynein light intermediate chain 1 (LIC1) (Tynan et al., 2000). Moreover, the LIC1-interacting domain on PCNT is mapped within ~580 amino acids located in the N-terminal half of PCNT (Tynan et al., 2000). Based on these findings, we propose a model in which the partially translated PCNT nascent polypeptide starts to interact with the dynein motor complex once the LIC1-interacting domain in the N-terminal half of PCNT is synthesized and folded, as early stages of protein folding can proceed quickly and co-translationally (Fedorov & Baldwin, 1997; Komar, Kommer, Krasheninnikov, & Spirin, 1997; Ptitsyn, 1995; Roder & Colon, 1997). Subsequently, this nascent polypeptide-dynein interaction allows the entire polysome, which is still actively translating *PCNT* mRNA, to be transported along the microtubule toward the centrosome (Figure 7). Alternatively, it is also possible that the coupling of the polysome to the motor complex is mediated through the ribosome-dynein interaction. If this was the case, additional components/adaptors would need to be involved in the interaction to differentiate the ribosomes translating *PCNT* mRNA from the ones translating other transcripts. One of the above mechanisms (i.e., via interaction through the nascent chain or ribosome itself) may also be used to mediate the co-translational transport of *ASPM* mRNA/polysomes to the mitotic spindle poles. Mapping the binding domains on both the motor and cargo sides, identifying the cargo adapter(s) that mediates the interaction, and testing the potential regulatory roles of mitotic kinases are important next steps to dissect the mechanisms underlying this co-translational protein targeting process.

**Figure 7.**
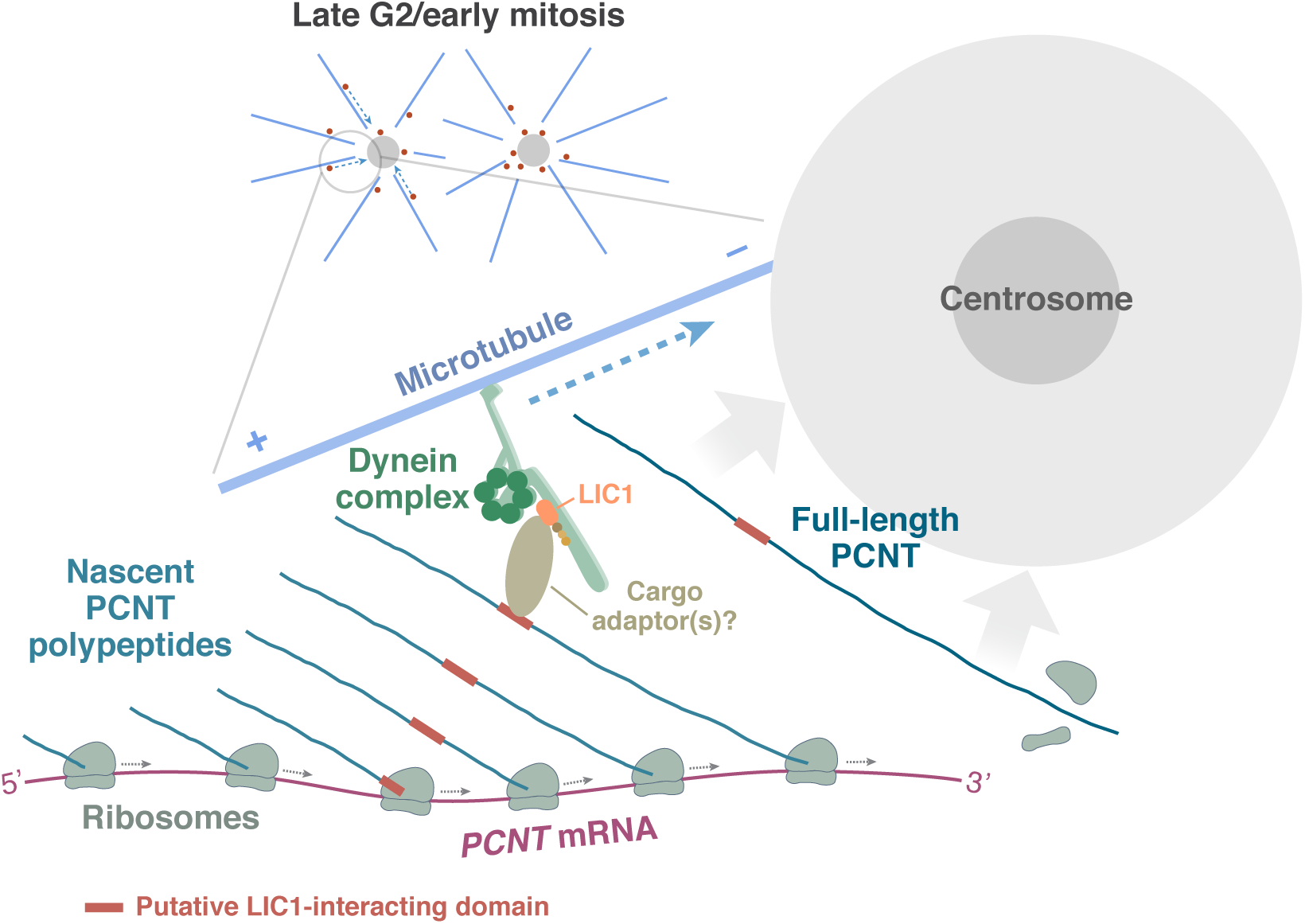
A model of co-translational targeting of PCNT polysomes to the centrosome during centrosome maturation. During the late G2/M transition, translation of *PCNT* mRNA is upregulated by an as yet unknown mechanism. The partially translated PCNT nascent polypeptide starts to interact with the dynein motor complex once the dynein light intermediate chain 1 (LIC1)-interacting domain in the N-terminal half of PCNT is synthesized and folded. Subsequently, this nascent polypeptide-dynein interaction allows the entire polysome, which is still actively translating *PCNT* mRNA, to be transported along the microtubule toward the centrosome. This co-translational targeting mechanism may maximize efficiency of PCNT production and delivery to the centrosome, prevent ectopic accumulation of PCNT outside of centrosomes, and/or facilitate integration of PCNT into the expanding PCM during early mitosis.

### Mitotic translation regulation of PCNT

Our data also underscore the importance of active translation of *PCNT* mRNA during early mitosis for the centrosome to gain the optimal level of PCNT because (1) during the G2/M transition, *PCNT* mRNA levels remain largely constant, but the centrosomal PCNT protein levels increase ~2-fold in 25 minutes after the onset of mitosis; (2) inhibiting translation briefly during early mitosis—e.g., 2 minutes of puromycin treatment in prophase or prometaphase—is sufficient to substantially reduce the amount of PCNT proteins incorporated at the centrosomes (Figure 5).

It is still unclear how the translation activation of *PCNT* mRNA is regulated during early mitosis. Previous studies show that translation is globally repressed during mitosis (Bonneau & Sonenberg, 1987; Fan & Penman, 1970; Pyronnet, Pradayrol, & Sonenberg, 2000) and this global translation repression is accompanied by the translation activation of a subset of transcripts through a cap-independent translation initiation mediated by internal ribosome entry sites (IRESes) (Cornelis et al., 2000; Marash et al., 2008; Pyronnet et al., 2000; Qin & Sarnow, 2004; Ramirez-Valle, Badura, Braunstein, Narasimhan, & Schneider, 2010; Schepens et al., 2007; Wilker et al., 2007). However, a recent study has challenged this view of IRES-dependent translation during mitosis and instead finds that canonical, cap-dependent translation still dominates in mitosis as in interphase (Shuda et al., 2015). Therefore, to elucidate the mechanism underlying the translation upregulation of *PCNT* mRNA during early mitosis, determining if this process is a cap- and/or IRES-dependent process might be a first logical step. In addition, our recent study has linked GLE1, a multifunctional regulator of DEAD-box RNA helicases, to the regulation of PCNT levels at the centrosome (Jao, Akef, & Wente, 2017). Since all known functions of GLE1 are to modulate the activities of DEAD-box helicases in mRNA export and translation (Alcazar-Roman, Tran, Guo, & Wente, 2006; Bolger, Folkmann, Tran, & Wente, 2008; Bolger & Wente, 2011; Weirich et al., 2006), it is worth elucidating whether translation upregulation of *PCNT* mRNA during mitosis is regulated through the role of GLE1 in modulating certain DEAD-box helicases involved in translation control such as DDX3 (H. H. Chen, Yu, & Tarn, 2016; Lai, Lee, & Tarn, 2008; Soto-Rifo et al., 2012).

In summary, the work presented here shows that incorporating PCNT into the PCM during centrosome maturation is at least in part mediated by upregulation of PCNT translation during the G2/M transition and the co-translational targeting of translating PCNT polysomes toward the centrosome during early mitosis. Efforts so far on elucidating the mechanism underlying centrosome maturation has focused for the most part on the interplay of protein-protein interactions and post-translational modifications (e.g., phosphorylation) of different PCM components. However, our study suggests that a spatiotemporal coupling between the active translation machinery and the motor-based transport may represent a new layer of control over centrosome maturation. Our work also suggests that spatially restricted mRNA localization and translation are not limited to early embryos or specialized cells (e.g., polarized cells such as neurons). We anticipate that co-translational protein targeting to subcellular compartments beyond the centrosome may prove to be a recurrent cellular strategy to synthesize and deliver certain cytoplasmic proteins to the right place at the right time. This regulatory process might represent an underappreciated, universal protein targeting mechanism, in parallel to the evolutionarily conserved co-translational targeting of secreted and membrane proteins to the ER for the secretory pathway.

## Materials and Methods

### Compounds

RO-3306 (4181, R&D Systems, Minneapolis, MN), ciliobrevin D (250401, MilliporeSigma, Burlington, MA), nocodazole (M1404, Sigma-Aldrich, St. Louis, MO), cytochalasin B (228090250, ACROS Organics, Geel, Belgium), emetine (324693, MilliporeSigma), puromycin (540222, MilliporeSigma), harringtonine (H0169, LKT Laboratories, St. Paul, MN).

### Antibodies

Antibodies were purchased commercially: rabbit anti-PCNT N-terminus (1:500 or 1:1,000 dilution, ab4448, Abcam, Cambridge, MA), anti-PCNT C-terminus (1:500 dilution, sc-28145, Santa Cruz Biotechnology Inc., Santa Cruz, CA), mouse anti-γ-tubulin (1:1,000 dilution, Clone GTU-88, T6557, Sigma-Aldrich), sheep anti-digoxigenin-alkaline phosphatase antibody (1:5,000 dilution, 11093274910, Roche Diagnostics, Mannheim, Germany), and sheep anti-digoxigenin-peroxidase antibody (1:500 dilution, 11207733910, Roche Diagnostics). Secondary antibodies were highly cross-adsorbed IgG (H+L) labeled with Alexa Fluor 488, 568, or 647 (1:500 dilution, all from Life Technologies, Carlsbad, CA).

### Zebrafish husbandry

Wild-type NHGRI-1 fish (LaFave, Varshney, Vemulapalli, Mullikin, & Burgess, 2014) were bred and maintained using standard procedures (Westerfield, 2000). Embryos were obtained by natural spawning and staged as described (Kimmel, Ballard, Kimmel, Ullmann, & Schilling, 1995). All animal research was approved by the Institutional Animal Care and Use Committee, Office of Animal Welfare Assurance, University of California, Davis.

### Generation of *pcnt* knockout fish

Disruption of zebrafish *pcnt* was generated by the CRISPR-Cas technology as described (Jao, Wente, & Chen, 2013). In brief, to generate guide RNA (gRNA) targeting *pcnt*, two complementary oligonucleotides (sequences in Supplementary Table 1) corresponding to a target sequence in the exon 2 of *pcnt* were annealed and cloned into pT7-gRNA plasmid to generate pT7-pcnt-gRNA. *pcnt* gRNA was generated by *in vitro* transcription using the MEGAshortscript T7 kit (AM1354, Thermo Fisher Scientific, Waltham, MA) with BamHI-linearized pT7-pcnt-gRNA as the template. Capped, zebrafish codon-optimized, double nuclear localization signal (nls)-tagged Cas9 RNA, *nls-zCas9-nls*, was synthesized by *in vitro* transcription using the mMESSAGE mMACHINE T3 kit (AM1348, Thermo Fisher Scientific) with XbaI-linearized pT3TS-nls-zCas9-nls plasmid as the template.

Microinjection of the mix of *pcnt* gRNA and *nls-zCas9-nls* RNA into zebrafish embryos (F0) was performed as described (Jao, Appel, & Wente, 2012). Pipettes were pulled on a micropipette puller (Model P-97, Sutter Instruments, Novato, CA). Injections were performed with an air injection apparatus (Pneumatic MPPI-2 Pressure Injector, Eugene, OR). Injected volume was calibrated with a microruler (typically ~1 nl of injection mix was injected per embryo). Injected F0 embryos were raised and crossed with wild-type zebrafish to generate F1 offspring. Mutations in F1 offspring were screened by PCR amplifying the target region (primer sequences are in Supplementary Table 2), followed by 7.5% acrylamide gel electrophoresis to detect heteroduplexes and sequencing. Two frameshift mutant alleles of *pcnt, pcnt^tup2^* and *pcnt^tup5^*, were used in this study (Figure 1-figure supplement 1). Maternal-zygotic *pcnt* mutant embryos were generated by intercrosses of homozygous *pcnt^tup2^* or *pcnt^tup5^* fish.

### Inhibition of protein synthesis of zebrafish early embryos

To inhibit protein synthesis in blastula-stage zebrafish embryos, one-cell stage embryos from wild-type NHGRI-1 intercrosses were injected with ~1 nl of Injection Buffer alone (10 mM HEPES, pH 7.0, 60 mM KCl, 3 mM MgCl_2_, and 0.05% phenol red) or with 300 μM puromycin in Injection Buffer. The embryos were fixed and analyzed after they developed to the 2-cell stage.

### Cell culture

HeLa cells (ATCC^®^ CCL-2™, a gift from Susan Wente, Vanderbilt University, Nashville, TN, or a HeLa cell line stably expressing scFv-sfGFP-GB1 and NLS-tdPCP-tdTomato, a gift from Xiaowei Zhuang, Howard Hughes Medical Institute, Harvard University, Cambridge, MA; Wang et al., 2016) and Centrin-GFP RPE-1 cells (a gift from Alexey Khodjakov, Wadsworth Center, New York State Department of Health, Rensselaer Polytechnic Institute, Albany, NY; Uetake et al., 2007) were maintained in Dulbecco’s Modification of Eagles Medium (10–017-CV, Corning, Tewksbury, MA) and Dulbecco’s Modification of Eagles Medium/Ham’s F-12 50/50 Mix (10–092-CV, Corning), respectively. All cell lines were supplemented with 10% fetal bovine serum (FBS) (12303C, lot no. 13G114, Sigma-Aldrich, St. Louis, MO), 1× Penicillin-Streptomycin (30–002 CI, Corning), and maintained in a humidified incubator with 5% CO_2_ at 37°C. To inhibit cytoplasmic dynein activities, the cells were treated with 50 μM ciliobrevin D for 1 hr 25 min at 37°C.

Cell lines used in this study were not further authenticated after obtaining from the sources. None of the cell lines used in this study were included in the list of commonly misidentified cell lines maintained by International Cell Line Authentication Committee.

### Cell synchronization

**Early M phase**. Cells were synchronized by either double thymidine block using 2 mM thymidine (J. Jackman & O'Connor, 2001) or by the RO-3306 protocol using 6 *μ*M RO-3306 (Vassilev et al., 2006). For HeLa and Centrin-GFP RPE-1 cells, prophase and prometaphase cells were enriched in the cell population ~8 hr after the second release in the double thymidine block protocol, or 20–25 min after releasing cells from an 18-hr RO-3306 treatment.

**G1 phase**. Cells were incubated with 6 *μ*M RO-3306 for 18 hr, washed out, and incubated in fresh media with 10% FBS for 30 min. Mitotic cells were collected after two firm slaps on the plate and were plated again to circular coverslips. The cells were grown for 6 hr; at this time, almost all cells are in G1 phase (i.e., two centrin dots per cell).

### RNA *in situ* hybridization in zebrafish

*In situ* hybridizations of zebrafish embryos were performed as described (Thisse & Thisse, 2008). In brief, the DNA templates for making *in situ* RNA probes were first generated by RT-PCR using Trizol extracted total RNA from wild-type zebrafish oocytes as the template and gene-specific primers with T7 or T3 promoter sequence (sequences in Supplementary Table 2). Digoxygenin-labeled antisense RNA probes were then generated by *in vitro* transcription and purified by ethanol precipitation (sequences in Supplementary File 1). Blastula-stage embryos were fixed 4% paraformaldehyde in 1 × PBS with 0.1% Tween 20 (1 × PBS-Tw) overnight at 4°C, manually dechorionated, and pre-hybridized in hybridization media (65% formamide, 5× SSC, 0.1% Tween-20, 50 *μ*g/ml heparin, 500 *μ*g/ml Type X tRNA, 9.2 mM citric acid for 2–5 hr at 70°C, and hybridized for ~18 hr with hybridization media containing diluted antisense probe at 70°C. After hybridization, embryos were successively washed with hybridization media, 2× SSC with 65% formamide, and 0.2× SSC at 70°C, and finally washed with 1× PBS-Tw at 25°C. Embryos were then incubated for 3–4 hr with blocking solution (2% sheep serum, 2 mg/ml BSA, 0.1% Tween-20, 1× PBS) at 25°C, and incubated ~18 hr with blocking buffer containing anti-digoxigenin-alkaline phosphatase antibody (1:5,000 dilution) at 4°C. Embryos were washed successively with 1 × PBS-Tw and AP Buffer (100 mM Tris, pH 9.5, 100 mM NaCl, 5 mM MgCl_2_, 0.1% Tween-20) before staining with the NBT/BCIP substrates (11383213001/11383221001, Roche Diagnostics) in AP Buffer.

For combined RNA *in situ* hybridization and immunofluorescence to label both the RNA and centrosomes in zebrafish embryos, the RNA *in situ* hybridization process was performed as described above until the antibody labeling step: The embryos were incubated for ~18 hr with blocking solution (2% sheep serum, 2 mg/ml BSA, 0.1% Tween-20, 1× PBS) containing anti-digoxigenin-peroxidase (1:500 dilution) and anti-γ-tubulin (1:1,000 dilution) antibodies at 4°C. Embryos were washed successively with 1× PBS-Tw and then incubated for ~18 hr with blocking solution containing Alexa Fluor 568 anti-mouse secondary antibody (1:500 dilution). After secondary antibody incubation, embryos were washed successively with 1 × PBS and borate buffer (37.5 mM NaCl, 100 mM boric acid, pH 8.5) with 0.1% Tween-20. The RNA was visualized after tyramide amplification reaction by incubating embryos for 25 min in tyramide reaction buffer (100 mM borate buffer, 37.5 mM NaCl, 2% dextran sulfate, 0.1% Tween-20, 0.003% H_2_O_2_, 0.15 mg/ml 4-iodophenol) containing diluted Alexa Fluor 488 tyramide at room temperature. The reaction was stopped by incubating embryos for 10 min with 100 mM glycine, pH 2.0 at room temperature, followed by successive washes with 1× PBS-Tw.

### Fluorescent *in situ* hybridization with tyramide signal amplification (TSA) in human cultured cells

In brief, the DNA templates for making *in situ* RNA probes were first generated by RT-PCR using Trizol extracted total RNA from human 293T cells as the template and gene-specific primers with T7 or T3 promoter sequence (sequences in Supplementary Table 2). Digoxygenin-labeled antisense RNA probes were then generated by *in vitro* transcription and purified by ethanol precipitation (sequences in Supplementary File 1). Cells were fixed for ~18 hr with 70% ethanol at 4°C, rehydrated with 2× SSC (0.3 M NaCl, 30 mM trisodium citrate, pH 7.0) containing 65% formamide at room temperature, pre-hybridized for 1 hr with hybridization media (65% formamide, 5× SSC, 0.1% Tween-20, 50 μg/ml heparin, 500 *μ*g/ml Type X tRNA, 9.2 mM citric acid) at 70°C, and hybridized for ~18 hr with hybridization media containing diluted antisense probes at 70°C. Cells were then successively washed with hybridization media, 2× SSC with 65% formamide, and 0.2× SSC at 70°C, and finally washed with 1× PBS at room temperature. For tyramide signal amplification, cells were washed with 1 × PBS, incubated for 20 min with 100 mM glycine, pH 2.0, and washed with 1× PBS at room temperature. Cells were then incubated for 1 hr with blocking buffer (2% sheep serum, 2 mg/ml BSA, 0.1% Tween-20, 1× PBS) at room temperature, and incubated ~18 hr with blocking buffer containing anti-digoxigenin-peroxidase antibody (1:500 dilution) at 4°C. Cells were washed successively with 1× PBS and borate buffer (37.5 mM NaCl, 100 mM boric acid, pH 8.5) with 0.1% Tween-20 and incubated for 5 min in tyramide reaction buffer (100 mM borate buffer, 37.5 mM NaCl, 2% dextran sulfate, 0.1% Tween-20, 0.003% H_2_O_2_, 0.15 mg/ml 4-iodophenol) containing diluted Alexa Fluor tyramide at room temperature. Cells were washed successively with 1 × quenching buffer (10 mM sodium ascorbate, 10 mM sodium azide, 5 mM Trolox, 1× PBS) and 1× PBS at room temperature. Coverslips were mounted using ProLong^®^ Antifade media (P7481, Life Technologies).

### Sequential immunofluorescence (IF) and RNA single molecule fluorescent *in situ* hybridization (smFISH)

Sequential IF and smFISH were performed according to the manufacturer’s protocol (LGC Biosearch Technologies, Petaluma, CA) with the following modifications: IF was performed first. Cells were fixed for 10 min in 4% paraformaldehyde in 1 × PBS, washed twice with 1 × PBS, and permeabilized with 0.1% Triton X-100 in 1× PBS for 5 min at room temperature. Cells were washed once with 1 × PBS and incubated with 70 *μ*l of diluted primary antibody in 1 × PBS for 1 hr at room temperature. Cells were washed three times with 1 × PBS and incubated with 70 *μ*l of diluted secondary antibody in 1 × PBS for 1 hr at room temperature. Cells were washed three times with 1× PBS and post-fixed for 10 min in 3.7% formaldehyde in 1× PBS at room temperature. For the smFISH process, cells were washed with Wash Buffer A, incubated with 67 *μ*l of Hybridization Buffer containing 125 nM DNA probes labeled with Quasar 670 (sequences in Supplementary File 1) for 6 hr at 37°C. Cells were then incubated with Wash Buffer A for 30 min at 37°C, Wash Buffer A containing 0.05 *μ*g/ml DAPI for 30 min at 37°C, and Wash Buffer B for 3 min at room temperature. Coverslips were mounted using ProLong^®^ Antifade media (Life Technologies) and sealed with clear nail polish before imaging.

### Immunofluorescence

Cells were fixed for 10 min in 4% paraformaldehyde in 1× PBS, washed twice with 1× PBS, and permeabilized with 0.5% Triton X-100 in 1× PBS for 5 min at room temperature. Cells were incubated with blocking solution (2% goat serum, 0.1% Triton X-100, and 10 mg/ml of bovine serum albumin in 1× PBS) for 1 hr at room temperature, incubated with blocking solution containing diluted primary antibody for 1 hr at room temperature. Cells were washed three times with 1 × PBS and incubated with blocking solution containing diluted secondary antibody for 1 hr at room temperature. Cells were washed with 1 × PBS and nuclei were counterstained with 0.05 *μ*g/ml of DAPI in 1× PBS for 20 min at room temperature before mounting.

### EdU labeling

S phase cells were detected by using the Click-iT™ EdU Imaging Kit (Life Technologies) according to the manufacturer’s instruction. In brief, Centrin-GFP RPE-1 cells were grown on 12-mm acid-washed coverslips and pulse labeled with 10 *μ*M 5-ethynyl-2’-deoxyuridine (EdU) for 30 min at 37°C. The cells were then fixed for 10 min with 4% paraformaldehyde in 1 × PBS at room temperature, washed twice with 1× PBS, and permeabilized for 20 min with 0.5% Triton X-100 in 1× PBS. Cells were then washed twice with 1× PBS and incubated with a Click-iT cocktail mixture containing Alexa Fluor^®^ 488 or 594 azide for 30 min in the dark at room temperature.

### Microscopy

Embryos subjected to *in situ* hybridization were mounted in a 35-mm glass bottom dish (P35G-1.5–10-C, MatTek, Ashland, MA) in 0.8% low melting point agarose and imaged using a stereo microscope (M165 FC, Leica, Wetzlar, Germany) with a Leica DFC7000 T digital camera.

Confocal microscopy was performed using either a Leica TCS SP8 laser-scanning confocal microscope system with 63×/1.40 or 100×/1.40 oil HC PL APO CS2 oil-immersion objectives and HyD detectors in resonant scanning mode, or a spinning disk confocal microscope system (Dragonfly, Andor Technology, Belfast, UK) housed within a wrap-around incubator (Okolab, Pozzuoli, Italy) with Leica 63×/1.40 or 100×/1.40 HC PL APO objectives and an iXon Ultra 888 EMCCD camera for smFISH and live cell imaging (Andor Technology). Deconvolution was performed using either the Huygens Professional (Scientific Volume Imaging b.v., Hilversum, Netherlands) (for images captured on Leica SP8) or the Fusion software (Andor Technology) (for images captured on Andor Dragonfly).

## Quantification of smFISH data and PCNT levels at centrosomes

To quantify the RNA distribution within the cell in 3D voxels, we used Imaris software (Bitplane, Belfast, UK) to fit the protein signal as surfaces and the mRNA signal as spots of different sizes in deconvolved images of each confocal z-stack. The intensity of the mRNA signal in each spot is assumed to be proportional to the amount of mRNA in each spot and is used in lieu of mRNA units. The outline of the cell was obtained either from a transmitted light image or from the background in the pre-deconvolved image and was used to restrict fitting of both mRNA and protein signals to the cell of interest. The distance from each mRNA spot to each centrosome’s center of mass was calculated and the mRNA signal was “assigned” to the closest centrosome. The mRNA spots were binned by distance to the centrosome and the intensities of the spots in each bin were added as a measure of the amount of mRNA at that distance. This was calculated for each cell and then averaged over all the cells for each condition. Thus, the graphs show average mRNA as a function of distance (binned in 0.5 *μ*m intervals).

To quantify PCNT intensities at the centrosome, we put the surfaces of the anti-PCNT signals fit on the deconvolved images over the original images and used the statistics function in Imaris (Bitplane) to obtain the intensity sum of the original images within the fit volume.

### Live translation assay (SunTag/PP7 system)

A HeLa cell line stably expressing scFv-sfGFP-GB1 and NLS-tdPCP-tdTomato was transfected with the SunTag/PP7 reporter plasmid pEF-24×V4-ODC-24×PP7 (Wang et al., 2016) using Lipofectamine 3000 transfection reagent (Life Technologies) according to the manufacturer’s instruction. 12–18 hr after transfection, the medium was changed to 10% FBS/DMEM without phenol red before imaging.

### Statistical analysis

Statistical analysis was performed using the GraphPad Prism 7. Each exact *n* value is indicated in the corresponding figure or figure legend. Significance was assessed by performing an unpaired two-sided Student’s t-test, as indicated in individual figures. The experiments were not randomized. The investigators were not blinded to allocation during experiments and outcome assessment.

## Author Contributions

Conceptualization, L.J.; Methodology, G.S., M.A., I.B., B.H., and L.J.; Software, I.B.; Formal Analysis, G.S., M.A., I.B., K.M., T.O., N.C., L.N.C., L.C., B.H., and L.J.; Investigation, G.S., M.A., I.B., K.M., T.O., N.C., L.N.C., L.C., D.Y., and L.J.; Resources, I.B. and L.J.; Writing - Original Draft, G.S., M.A., I.B., and L.J.; Writing - Review & Editing, G.S., M.A., I.B., and L.J.; Visualization, I.B. and L.J.; Supervision, L.J.; Project administration, L.J.; Funding acquisition, L.J.

## Acknowledgement

We thank Susan Wente for the HeLa cell line; Alexey Khodjakov for the Centrin-GFP RPE-1 stable cell line; Xiaowei Zhuang for the SunTag/PP7 reporter plasmid pEF-24×V4-ODC-24×PP7, and the HeLa cell line stably expressing scFv-sfGFP-GB1 and NLS-tdPCP-tdTomato; Dena Leerberg and Bruce Draper for technical help on fluorescent *in situ* hybridization in zebrafish; Tom Glaser, Henry Ho, Frank McNally, Richard Tucker, and Mark Winey for critical reading of the manuscript; Emily Jao for help on digital illustrations. Experiments were performed in part through the use of UC Davis Health Sciences District Advanced Imaging Facility.

---

**Figure 1-figure supplement 1.**
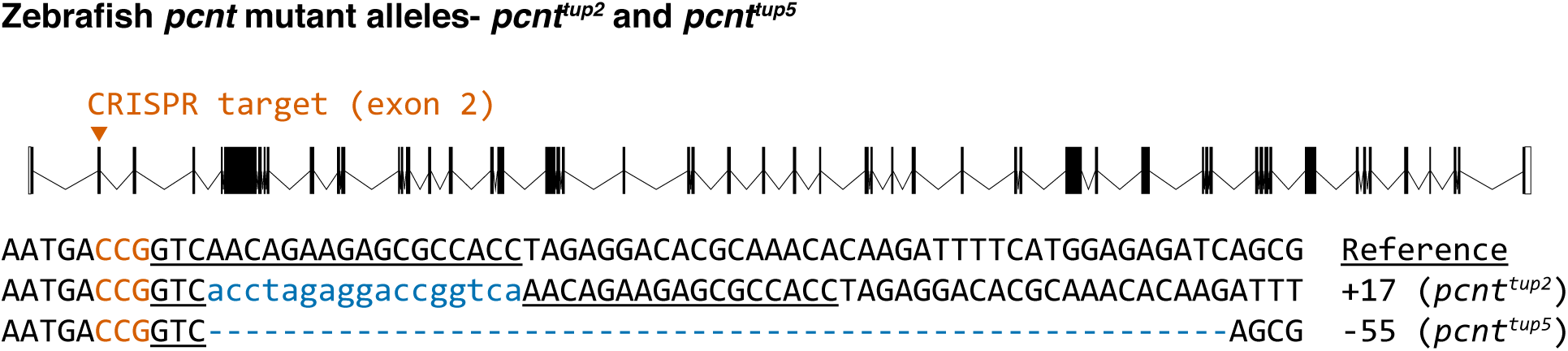
Sequences of two Cas9-induced frameshift mutations (alleles *pcnt^tup2^* and *pcnt^tup^*^5^) in the zebrafish *pcnt* gene. The wild-type reference sequence is on the top. The guide RNA targets the exon 2 of the *pcnt* transcript (encoded by ENSDARG00000033012). The target site is underlined and the proto-spacer-adjacent motif (PAM) is in orange (on the reverse strand). Insertions and deletions (indels) are indicated by blue lowercase letters and dashes, respectively. The net change of each indel mutation is noted at the right of each sequence (+, insertion; −, deletion).

The following figure supplements are available for **Figure 2:**

**Figure 2- figure supplement 1.**
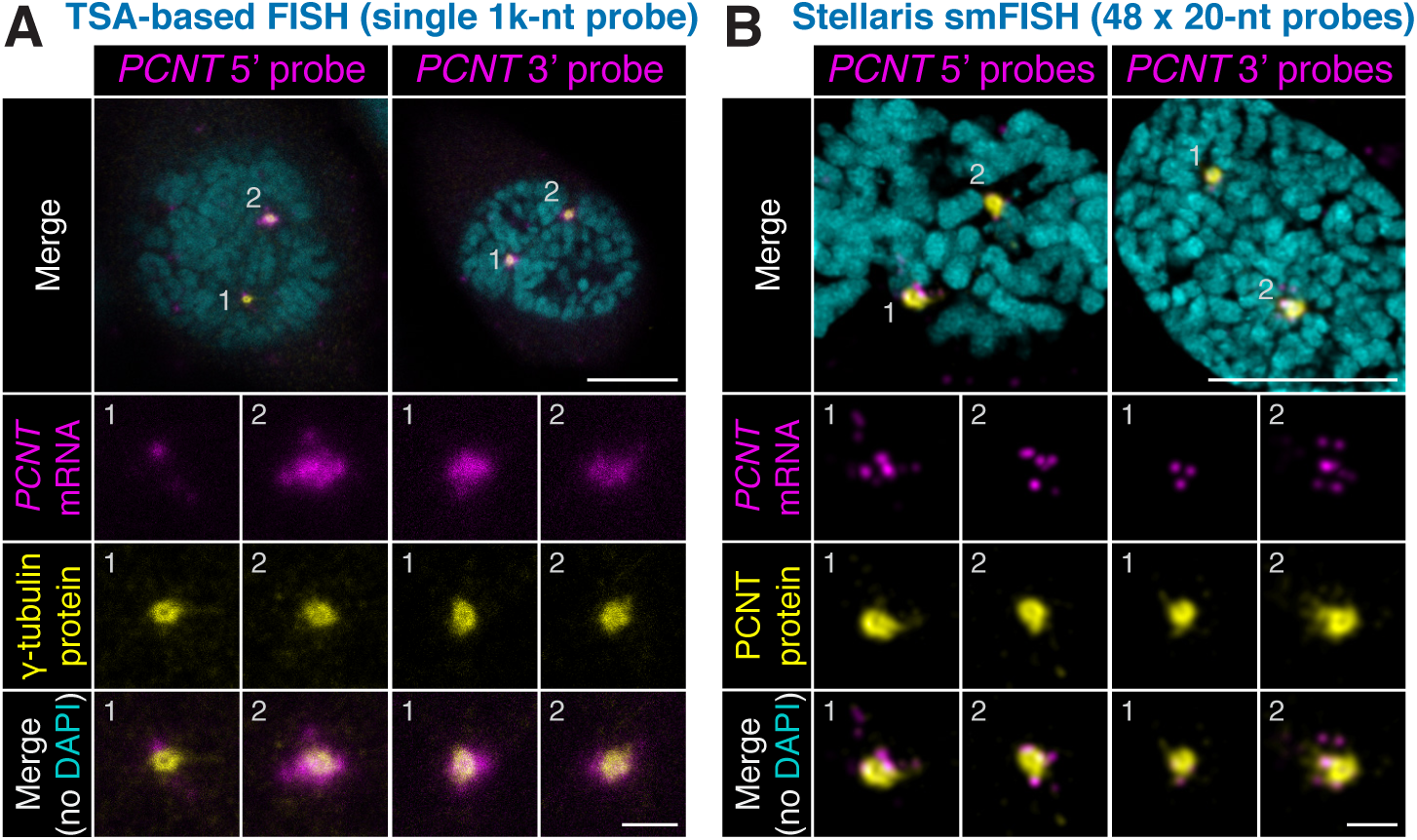
Non-overlapping antisense probes and two independent *in situ* methods confirm centrosomal localization of *PCNT* mRNA during early mitosis. HeLa cells were subjected to fluorescent *in situ* hybridization (FISH) with tyramide signal amplification (TSA) against *PCNT* mRNA with a 1,000-nt probe **(A)** or the Stellaris single-molecule FISH (smFISH) with a set of 48 20-nt fluorescent probes **(B)**. Note that both methods (probing two distinct regions in each method) showed similar distributions of *PCNT* mRNA at centrosomes during early mitosis, with the Stellaris smFISH showing mRNA at near single-molecule resolution. Scale bars: 10 *μ*m and 2 *μ*m (inset).

**Figure 2- figure supplement 2.**
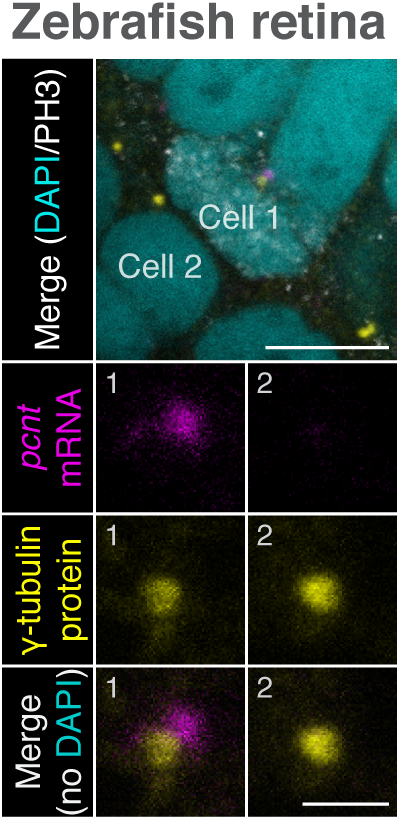
Zebrafish *pcnt* mRNA is localized to centrosomes of mitotic retinal neuroepithelial cells *in vivo*. Retinal neuroepithelial cells of 1 day old zebrafish were subjected to *pcnt* FISH, anti-γ-tubulin, and anti-phospho-Histone H3 (PH3) immunostaining. Note that *pcnt* mRNA was localized to the centrosome of the mitotic (Cell 1, PH3^+^), but not of the non-mitotic cell (Cell 2). Scale bars: 10 *μ*m and 2 *μ*m (inset).

**Figure 3- figure supplement 1.**
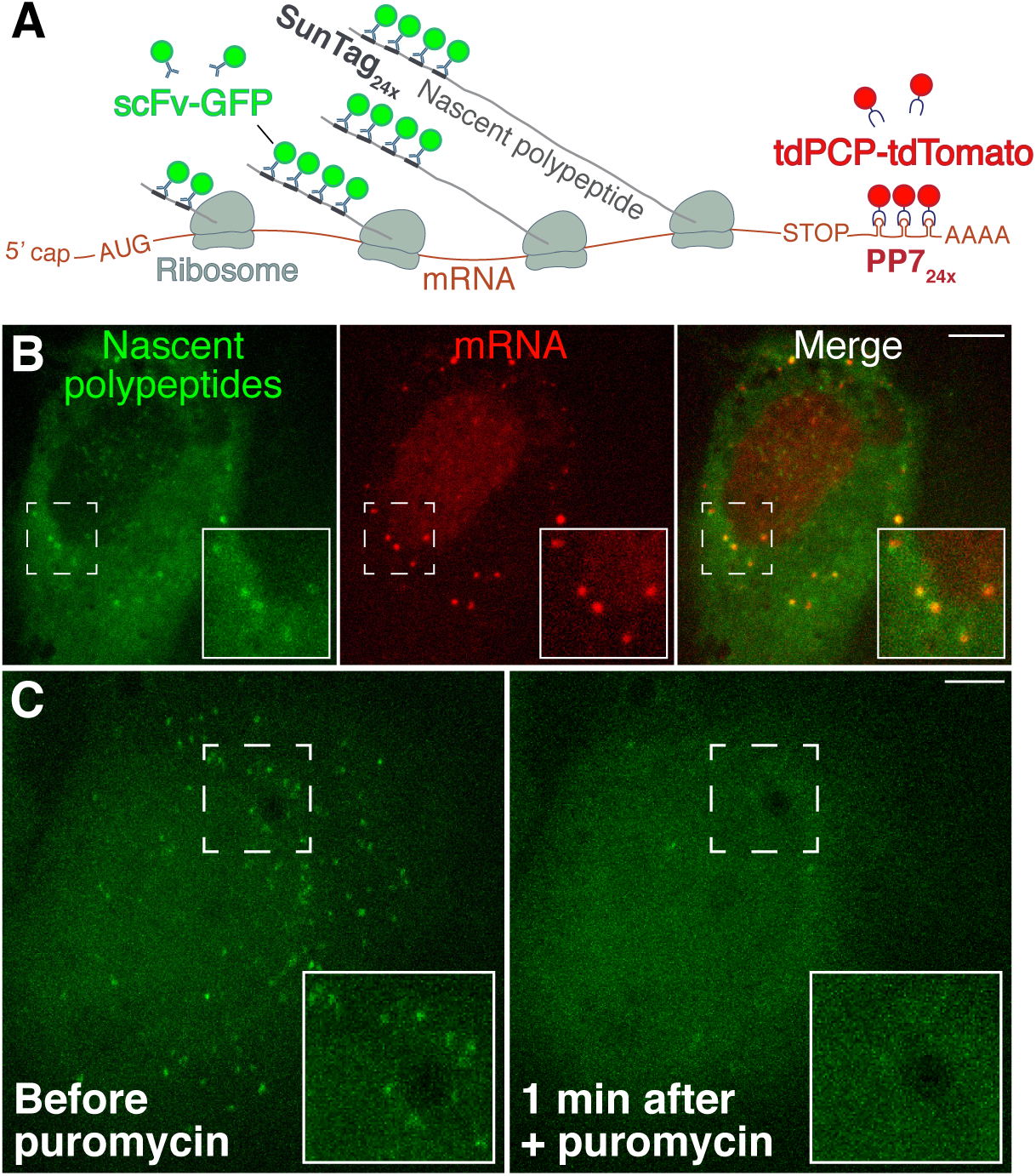
Visualization of active translation in live cells using the SunTag/PP7 system. **(A)** SunTag/PP7 system overview, adapted from Wang et al., 2016. **(B)** HeLa cells stably expressing scFv-GFP and tdPCP-tdTomato transfected with SunTag-ODC-PP7 reporter. Individual polysomes (GFP^+^) and mRNA (tdTomato^+^) were shown. **(C)** Translation foci in the same field before and after adding 300 *μ*m puromycin for 1 minute. Scale bars: 10 *μ*m.

**Figure 3- figure supplement 2.**
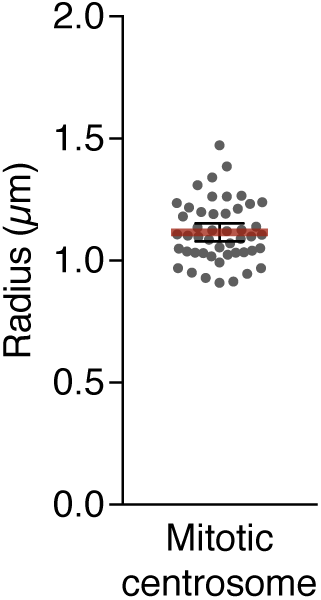
Mean radius of mitotic centrosomes of HeLa cells. Data are represented as mean ± 95% Cl.

**Figure 4- figure supplement 1.**
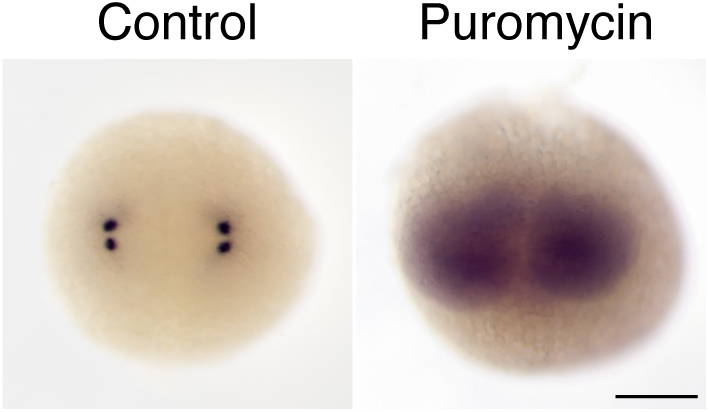
Centrosomal localization of zebrafish *pcnt* mRNA depends on intact polysomes. RNA *in situ* hybridization showed that *pcnt* transcripts were localized to the centrosomes in the buffer-injected embryo (Control), but were diffused throughout the cell in the embryo injected with ~1 nl of 300 *μ*m puromy-cin at the 1-cell stage (Puromycin). Both embryos shown are at the 2-cell stage. Scale bar: 200 *μ*m.

**Figure 4- figure supplement 2.**
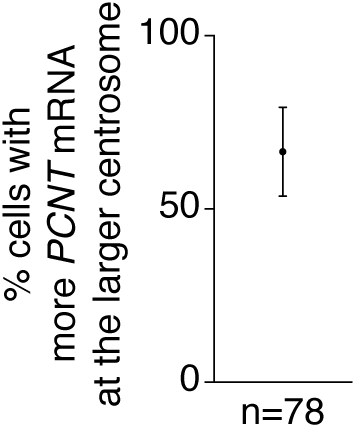
More *PCNT* mRNA was often enriched near the larger centrosome in early mitosis. In the majority of pro- and prometaphase HeLa cells (~67%), more *PCNT* mRNA was enriched around the larger centrosome. Data are represented as mean ± SD, “n” indicates the total number of cells analyzed from four experiments.

**Figure 4- figure supplement 3.**
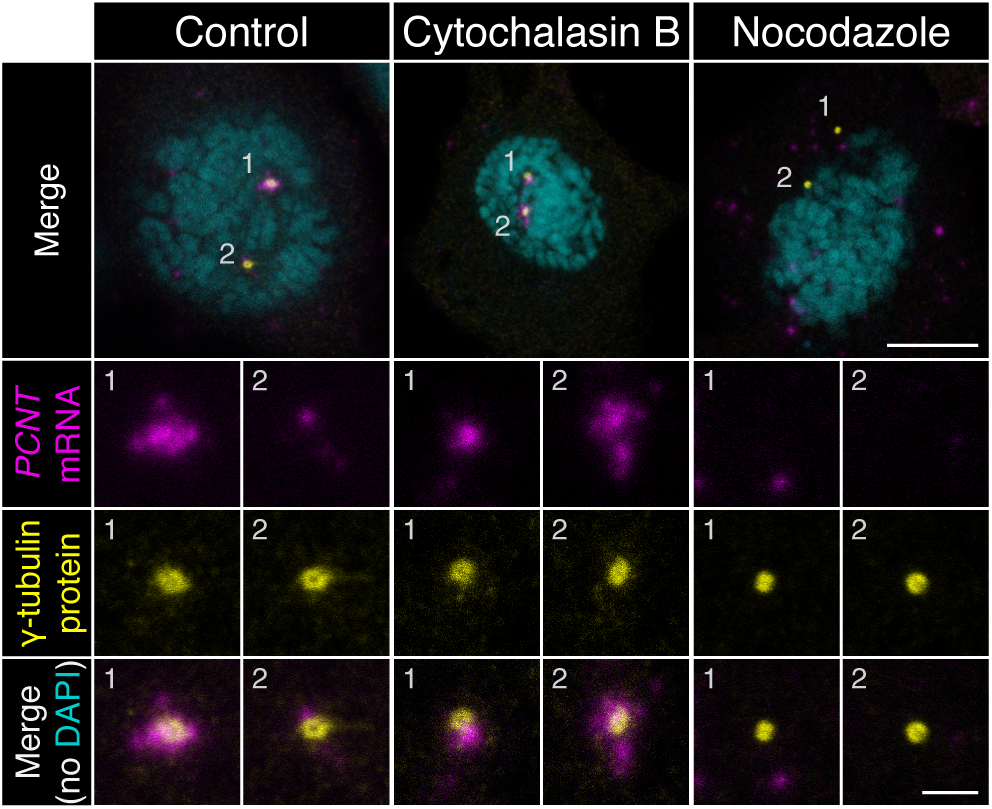
Centrosomal localization of human *PCNT* mRNA during early mitosis is microtubule-dependent. HeLa cells synchronized at prophase (pro) were treated with 5 *μ*g/ml cytochalasin B for 15 minutes or 10 *μ*g/ml nocodazole for 30 minutes at 37°C before fluorescent *in situ* hybridization with tyramide signal amplification against *PCNT* mRNA and anti-γ-tubulin immunostaining. Note that nocodazole, but not cytochalasin B, disrupted the centrosomal enrichment of *PCNT* mRNA. Scale bars, 10 *μ*m and 2 *μ*m (inset).

